# Mechanical forces impair antigen discrimination by reducing differences in T cell receptor off-rates

**DOI:** 10.1101/2022.05.05.490751

**Authors:** Johannes Pettmann, Lama Awada, Bartosz Różycki, Anna Huhn, Sara Faour, Mikhail Kutuzov, Laurent Limozin, Thomas R. Weikl, P. Anton van der Merwe, Philippe Robert, Omer Dushek

## Abstract

T cells use their T cell receptors (TCRs) to discriminate between lower-affinity self and higher-affinity foreign peptide major-histocompatibility-complexes (pMHCs) based on the TCR/pMHC off-rate. It is now appreciated that T cells generate mechanical forces during this process but how force impacts the TCR/pMHC off-rate remains unclear. Here, we measured the effect of mechanical force on the off-rate of multiple TCR/pMHC interactions. Unexpectedly, we found that lower-affinity pMHCs with faster solution off-rates were more resistant to mechanical force (weak slip or catch bonds) than higher-affinity interactions (strong slip bonds), and this was confirmed by molecular dynamic simulations. Consistent with these findings, we show that the best characterized catch-bond, involving the OT-I TCR, has a low affinity and an exceptionally fast solution off-rate. Our findings imply that reducing forces on the TCR/pMHC interaction improves antigen discrimination and we suggest this new force-shielding role for the adhesion receptors CD2 and LFA-1.

**One sentence summary:** Mechanical forces disproportionately accelerate the off-rates of higher-affinity antigens reducing T cell antigen discrimination

## Introduction

T cells use their T cell antigen receptors (TCRs) to recognise peptide antigens bound to major-histocompatibility-complexes (pMHCs), on the surface of antigen-presenting-cells (APCs). It is now well established that T cells discriminate between self and foreign pathogen (or cancer) derived pMHC based on the kinetic off-rate of the interaction (1–4). Consistent with this mechanism, the off-rate (*k*_off_) measured in solution with purified TCR and pMHC can usually predict the T cell response. Recently, a number of studies have shown that T cells can generate forces as they probe surfaces for pMHC (5–7) and can impose forces directly on TCR/pMHC interactions (7, 8). This observation is important because T cells discriminate antigens based on the *k*_off_ in the membrane, termed the membrane off-rate or 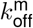 (Figure 1A), and this can be affected by force (9). Precisely how molecular forces on the TCR/pMHC interaction impact the 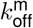 and antigen discrimination is controversial.

**Figure 1.**
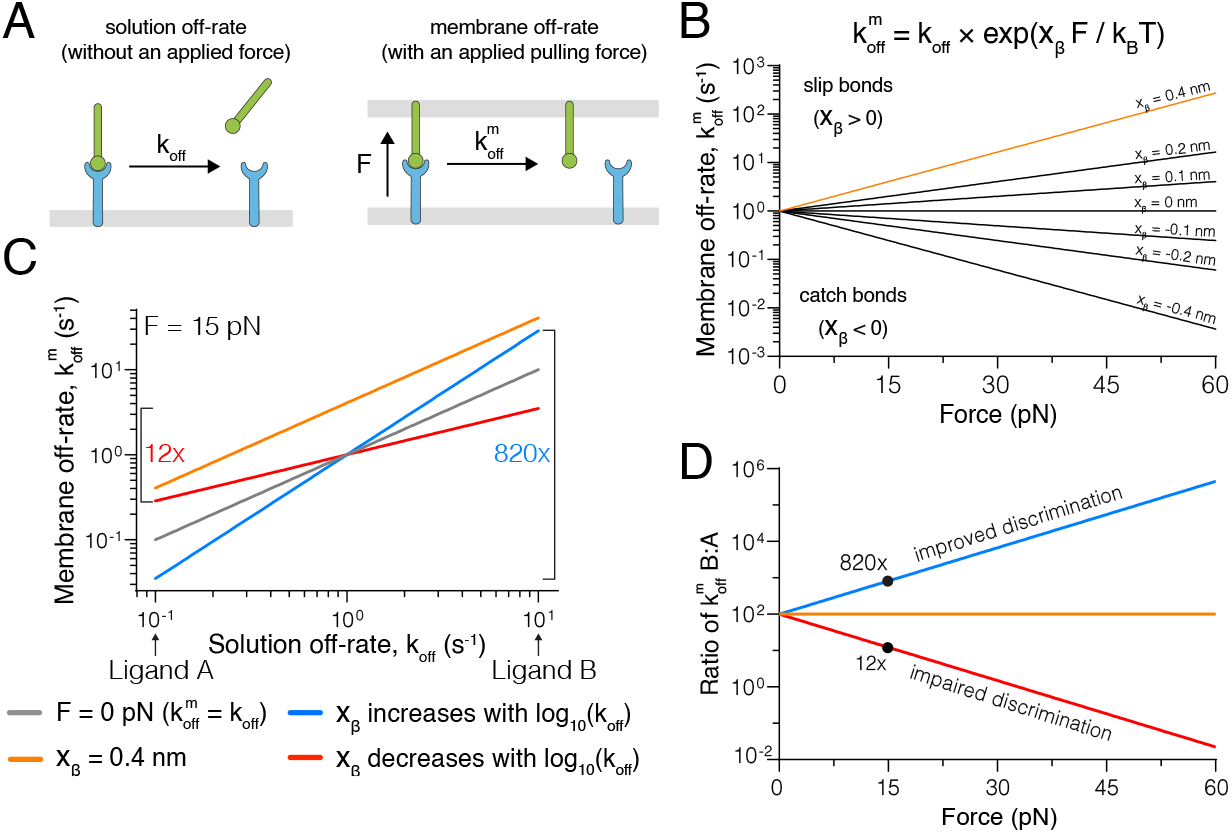
Modeling predicts that force can improve or impair antigen discrimination. **(A)** Depiction of (left) the dissociation of a soluble ligand and its solution off-rate (*k*_off_) and (right) a membrane-anchored ligand and its membrane off-rate 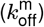, where the interaction is exposed to a pulling force (F). **(B)** The dependence of 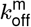 on the pulling force at the indicated force sensitivity parameters (x_β_) with *k*_off_ = 1 s^*−*1^. **(C)** Dependence of 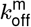on *k*_off_ at zero force (gray) or under an applied force of 15 pN (coloured lines), when x_β_ is constant (orange), positively (blue), or negatively (red) correlated with 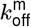In this example a 100-fold change in *k*_off_ (Ligand A versus Ligand B) can be increased to a *∼*820-fold or decreased to a *∼*12-fold change in 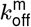, depending on whether x_β_ is positively or negatively correlated with the *k*_off_, respectively. **(D)** The fold-change in 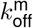 for Ligand B versus Ligand A over the applied force. All calculations are performed using the formula in (B) with *x*_*β*_ = 0.4 nm (orange), *x*_*β*_ = +0.3log_10_(*k*_off_) (blue), and *x*_*β*_ = *−*0.3log_10_(*k*_off_) (red) in panels C and D.

One hypothesis is that molecular force improves discrimination between low affinity self and high affinity foreign antigens (9). This is based on the assumption that force increases the 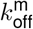 of low affinity pMHC interactions (termed slip-bonds) while decreasing the 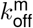 of higher affinity pMHC interaction (catch-bonds), thus magnifying differences in *k*_off_. Evidence for catch-bonds has been obtained mainly using the bio-force probe (10–13), which applies external forces to native TCRs on T cells. It is notable that the magnitude of these catch-bonds is appreciably reduced (14) or abolished (15) when applying force to purified forms of the same TCRs. This suggests that some catch bonds observed with the bioforce probe may be secondary to TCR signalling rather than intrinsic to the TCR/pMHC interaction. Further studies of a wider range of TCR/pMHC interactions are required to elucidate the relationship between force, kinetics, and T cell activation.

Several mechanisms are likely to influence the forces experienced by TCR/pMHC interactions. Firstly, in order for TCRs to engage pMHC the T cell and APC membranes need to be brought to within *∼* 14 nm (16), which requires compression of the large molecules that form the glycocalyx, such as CD45 and CD43, which span 21-45 nm (17, 18). By resisting compression, these glycocalyx molecules would generate forces on the TCR/pMHC interaction that are predicted to be *∼* 20 pN (19). Secondly, the dynamics of actin-based microvilli-like protrusions that form close contacts between T cells and APC membranes may directly or indirectly pull on TCRs (20–22). Finally, adhesion receptor/ligand interactions are likely to modulate the forces experiences by TCR/pMHC interactions (23). For example, the adhesion receptors CD2 and LFA-1 can improve antigen sensitivity (18) and antigen discrimination (4), and it is plausible that they do so, at least in part, by influencing the forces experienced by TCR/pMHC interactions.

Here, we found that force increased the *k*_off_ of most TCR/pMHC interactions. Unexpectedly, lower-affinity interaction were least sensitive to force and more likely to form catch bonds than high affinity interactions, which showed the highest sensitivity to force and this was also observed with molecular dynamic simulations. We show that the best characterized catch bond, involving the OT-I TCR, has an unusually low affinity and the fastest *k*_off_ yet reported for an agonist TCR/pMHC interaction. Our results imply that force will reduce differences between TCR/pMHC 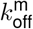 and therefore impair antigen discrimination. By incorporating force into the kinetic proof-reading model, we show that force-shielding can account for the ability of adhesion receptors to enhance antigen discrimination. Our study clarifies the role of force in T cell antigen recognition and reconciles apparently contradictory reports.

## Results

### Theoretical modeling predicts that mechanical forces can improve or impair antigen discrimination

To investigate how forces affect antigen discrimination, we used Bell’s well-established phenomenological model (24). It provides a simple relationship between the applied force and 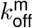, which depends on a receptor/ligand force sensitivity parameter (xβ). When xβ*>* 0, force *increases* 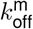 forming a slip-bond and when x_β_*<* 0, force *decreases* 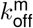 forming a catch-bond (Figure 1B). In short, the sensitivity of the bond to force increases as the value of its force sensitivity parameter increases.

If all pMHCs that bind the same TCR had the same force sensitivity (i.e. x_β_ is constant), then an applied force will increase the *k*_off_ of all pMHCs by the same factor (Figure 1C, orange). Because of this, the ratio of the 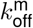 between any two pMHC ligands will be be unaffected by force (Figure 1D). Given that antigen discrimination is dependent on the fold change in 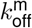 (9, 16), it follows that, if force sensitivity is constant, an applied force would not impact antigen discrimination.

We next explored how force would affect antigen discrimination if the force sensitivity is not constant but varies with *k*_off_ (or affinity, if *k*_on_ remains constant). Indeed, it has been proposed that higher-affinity TCR ligands form catch bonds whereas lower-affinity ligands form slip-bonds (10, 25), implying that the force sensitivity increases as the *k*_off_ increases. We confirmed this using our model, which shows that applied force amplifies the fold change in *k*_off_ to produce larger fold changes in 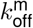 (Figure 1C,D blue). Conversely, if the force sensitivity decreases as the *k*_off_ increases then applied force dampens differences in *k*_off_ producing smaller fold changes in 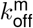 (Figure 1C,D red). This is illustrated with two test ligands that differ in *k*_off_ by 100-fold showing that the fold-change in 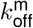 is *>*100-fold when x_β_ increases with *k*_off_ (i.e. improved discrimination) but *<*100-fold when x_β_ decreases with *k*_off_ (i.e. impaired discrimination) under an applied force (Figure 1D).

### The force-sensitivity of TCR/pMHC interactions decreases with off-rate or dissociation constant

Given that the impact of force on antigen discrimination critically depends on the relationship between *k*_off_ and x_β_, we set out to measure this using our laminar flow chamber apparatus (15, 26). In this assay, beads coated with TCR flow over a low density of pMHC while a camera records bead arrests (Figure 2A). The force exerted on the TCR/pMHC bond increases with flow velocity (26). Bead arrests are confirmed to be mediated by single bonds by showing that (1) the binding linear density (BLD) of the beads increases in proportion to the concentration of pMHC on the surface (Figure S1A) and that (2) the survival curves were independent of pMHC density (Figure S1B) (15, 26, 27).

**Figure 2.**
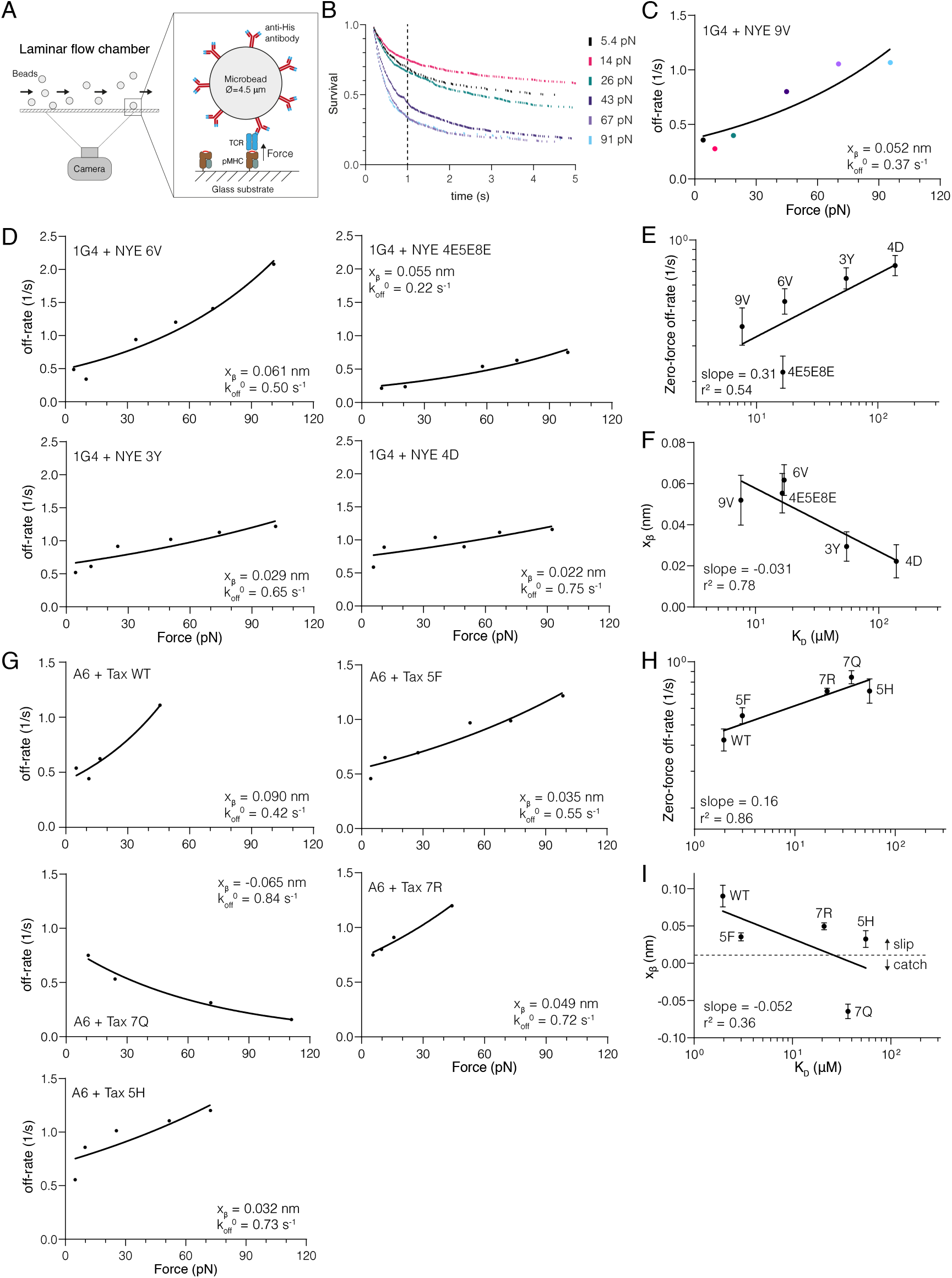
Direct measurements of force sensitivity (*x*_*β*_) for the 1G4 and A6 TCRs reveal a negative correlation with K_D_ (*k*_off_/*k*_on_). (A) The laminar flow chamber assay. All experiments were performed at physiological temperatures (37°C) and forces (*<*100 pN). (B) Example of bead survival distributions for the 1G4 TCR binding NYE 9V. The survival probability at 1 s is shown as a dotted vertical line and is used to calculate the off-rate under force: *−* ln(survival at 1 s)*/*1*s*. **(C)** Off-rates under force obtained from (B) are fitted with Bell’s model (solid line). **(D)** Off-rates under force for additional pMHCs interacting with the 1G4 TCR fitted to Bell’s model (solid line). **(E)** Correlation of the dissociation constant (K_D_) measured previously by SPR and the extrapolated zero-force flow chamber off-rate 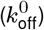 from fitting Bell’s model for the 1G4 TCR. **(F)** Correlation of the SPR dissociation constant (K_D_) with the fitted force sensitivity parameter (x_β_) for the 1G4 TCR. **(G)** Off-rates under force for pMHCs interacting with the A6 TCR fitted to Bell’s model (solid line). **(H)** Correlation of the dissociation constant (K_D_) measured previously by SPR and the extrapolated zero-force flow chamber off-rate 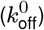 from fitting Bell’s model for the A6 TCR. **(I)** Correlation of the SPR dissociation constant (K_D_) with the fitted force sensitivity parameter (x_β_) for the A6 TCR. Error bars (SE) are obtained from the fit. The number of independent experiments that were combined to produce the estimated off-rates are: 7 (1G4/9V), 8 (1G4/6V), 8 (1G4/4E5E8E), 10 (1G4/3Y), 6 (1G4/4D), 9 (A6/Tax WT), 11 (A6/5F), 9 (A6/7Q), 8 (A6/7R), 8 (A6/5H)

We recently obtained accurate solution affinities for the 1G4 and A6 TCRs binding a panel of peptide ligands presented on HLA-A*02:01 (4), and so we selected several pMHCs from this panel (Figure S2). Survival distributions were generated using the arrest duration of individual beads; increasing force decreased survival for 1G4 binding the agonist 9V peptide from the NY-ESO-1 cancer testis antigen, which is indicative of a slip bond (Figure 2B). We calculated off-rates based on the survival fraction at 1 s as in our previous work (26) and fit the data using Bell’s model to determine the force sensitive and the zero-force off-rate (Figure 2C). The zero-force off-rate 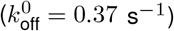 agreed with the published solution *k*_off_ obtained using SPR (0.33 s^*−*1^) (1). We found that four additional ligands also formed slip bonds (Figure 2D) and as expected, the extrapolated zero-force off-rate correlated with the K_D_ values measured by SPR (Figure 2E). We used K_D_ values because they correlate with the solution off-rates and are easier to measure accurately, especially for weakly binding pMHCs, where the solution off-rates are often too fast to measure.

We noted that the force sensitivity x_β_ varied between the pMHCs (Figure 2C,D) and, surprisingly, displayed a negative correlation with their K_D_ (Figure 2F). This means that an applied force disproportionately accelerates the off-rate of higher-affinity interactions. This can be observed by comparing the slopes on the off-rate curves (Figure 2D) where the higher-affinity 6V pMHC (top left panel) shows a much larger increase in off-rate with applied force than the lower affinity 4D pMHC (bottom right panel).

We next investigated whether the negative correlation could be observed for a different TCR. We found that the A6 TCR formed a slip bond with its wild-type Tax peptide from Human T-lymphotropic virus (HTLV), slip-bonds with three additional ligands (5F, 7R, 5H), and one catch bond (7Q) (Figure 2G). The estimated zero-force off-rate correlated with K_D_ measured by SPR (Figure 2H). As seen with the 1G4 TCR, we observed a negative correlation between the force sensitivity x_β_ and the K_D_ (Figure 2I).

We found three interactions (the 1G4 TCR binding 3A and 6T, and the A6 TCR binding 7T) that did not display canonical slip or catch bonds and could not be accurately fit with Bell’s model (Figure S3). These interactions displayed the expected binding linear densities (Figure S1). Importantly, these bonds were slip bonds at the physiologically relevant forces (≲20pN). Interestingly, 3A and 6T did display canonical slip and catch bonds, respectively, at 25°C (26), suggesting that some TCR/pMHC interactions can have a more complex unbinding pathway at 37°C.

We previously measured force sensitivity for the 1G4 TCR at 25°C using larger forces (*>*30 pN) (26). In that study, which used a different definition of the force sensitivity parameter (*F*_0_ = *k*_B_*T/x*_*β*_), it was reported that no significant (linear) correlation was observed between between *k*_off_ and *F*_0_. However, when we plot our force sensitivity parameter x_*β*_ against the log of *k*_off_, we observed a striking negative correlation (Figure S4A). Therefore, our previous result are consistent with the present study.

We identified only one other study that measured force sensitivity and affinity. In that study *x*_*β*_ was measured for a panel of nine antibodies binding fluorescein using atomic force microscopy (28). Interestingly, a striking negative correlation is also observed between force sensitivity and off-rate, which we reproduce by directly plotting *x*_*β*_ over *k*_off_ (Figure S4B). This suggests that a negative correlation between force sensitivity and off-rate (or K_D_) may be a general feature of antigen receptor interactions.

### Molecular dynamic simulation predicts a negative correlation between force sensitivity and off rate

We next determined whether the negative correlation between *k*_off_ and x_β_ can also be observed using molecular dynamics simulations. We employed a coarse-grained model (29) where the C-terminus of the TCRβ was fixed in space and the C-terminus of the MHC was moved with constant speed (*ν*) to generate a response force (*F*) on the TCR/pMHC interaction. Unbinding could readily be observed by *F* dropping from its maximal value to zero (Figure 3A). The maximum force (*F*_max_) was determined from independent simulations (Figure 3B) and plotted over *ν* (Figure 3C). Fitting the Bell-Evans formula (equation 2 in (30)) provides estimates of *k*_off_*τ* and x_β_, where *τ* is the simulation timestep (*∼*1 ns) and is constant for all calculations.

**Figure 3.**
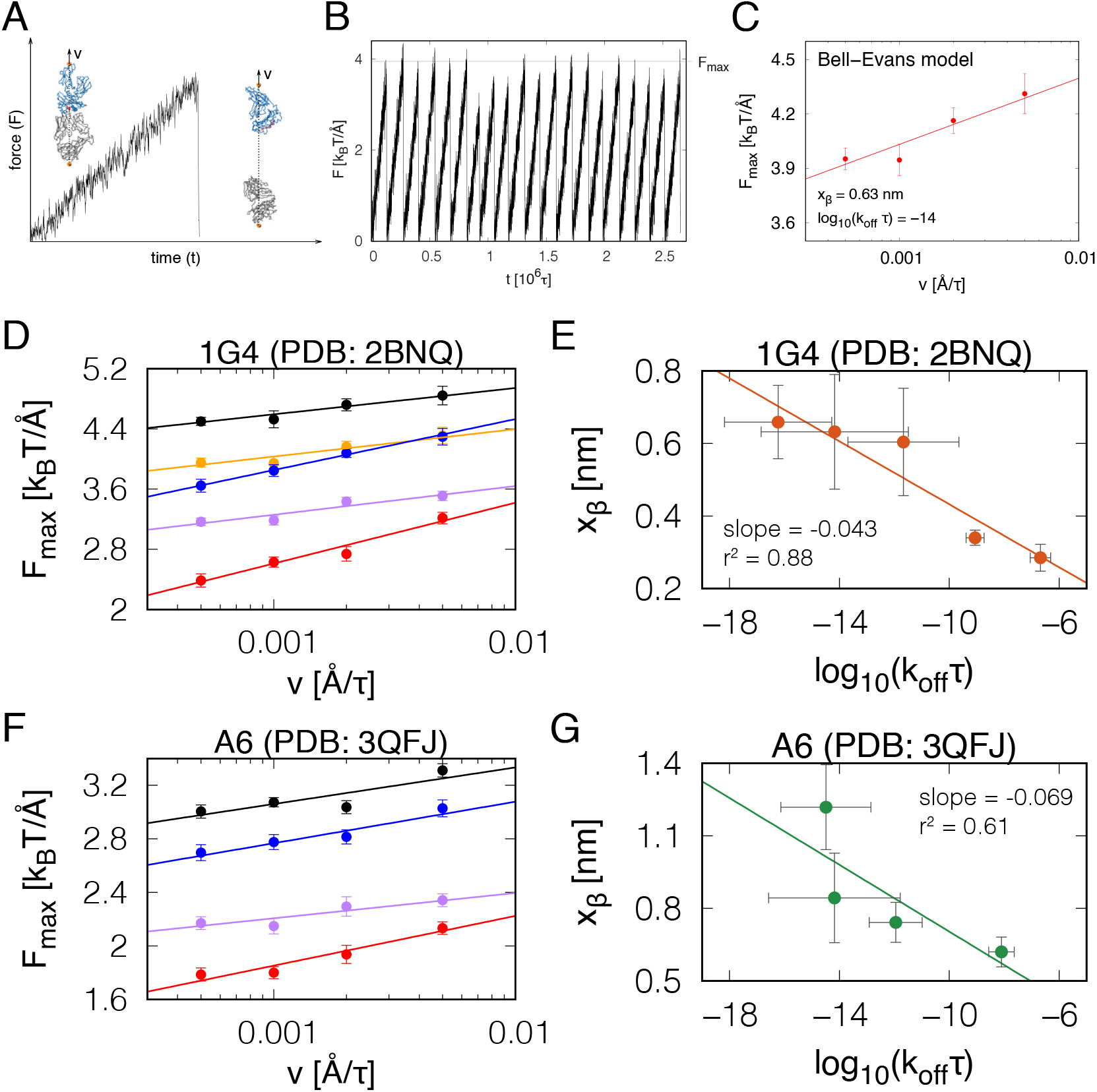
Coarse-grained molecular dynamics simulations support a negative correlation between x_β_ and *k*_off_. **(A)** The response force (*F*) on the TCR/pMHC interaction when pulled at acting on the spring attached to the C-terminus of the MHC which is being pulled with constant velocity (*ν*). A second spring is attached to the C-terminus of the TCRβ and fixed in space. Insets: the TCR is shown in grey, the MHC in blue, and the peptide in red. The sites where the springs are attached are marked in orange. The direction of pulling is marked by a dashed line. **(B)** A plot of *F* (*t*) from twenty independent simulation trajectories with the same value of the pulling speed (*ν*). The grey horizontal line indicates the average force (*F*_max_) at which the TCR-pMHC complex dissociates. **(C)** The dependence of *F*_max_ (N=20) on *ν* from the simulations follows the Bell-Evans model (solid line), which produces estimates of *k*_off_ and x_β_. **(D)** The dependence of *F*_max_ on *ν* obtained from simulations of five complexes for the 1G4 TCR-pMHC complex (PDB: 2BNQ): (i) with all native contacts between the peptide and the TCR (black), (ii) without the native contacts between the peptide residues 4 to 8 and the TCR (orange), (iii) only with the native contacts between the peptide and the TCRα (blue), (iv) only with the native contacts between the peptide and the TCRβ (purple), and (v) with no native contacts between the peptide and the TCR (red) included in the coarse-grained model. Solid lines are the fit of the Bell-Evans model. **(E)** The values of *k*_off_ and x_β_ from the fit of the Bell-Evans model to the simulation data shown in panel D. The solid line is a linear fit on log-transformed x-axis values. **(F)** Analogous to panel D but for the A6 TCR-pMHC complex (PDB: 3QFJ). The colour code is as in panel D. **(G)** The values of *k*_off_ and x_β_ from the fit of the Bell-Evans model to the simulation data shown in panel F. The solid line is a linear fit on log-transformed x-axis values.

To generate TCR/pMHC complexes with different affinities, we performed simulations using the 1G4 TCR/pMHC complex where all native contacts between the peptide and TCR were included or where certain contacts were excluded (Figure 3D). We fitted the Bell-Evans model and consistent with our experimental data, a plot of x_β_ over *k*_off_ from these simulations revealed a negative correlation with a dimensionless slope of -0.043 (Figure 3E), which is similar to the experimental slope of -0.031 observed for 1G4 (Figure 2F). We repeated the analysis for A6 TCR/pMHC complexes (Figure 3F) also finding a negative correlation with a dimensionless slope of -0.069 (Figure 3G), which again was similar to the experimental slope of -0.052 (Figure 2I). Lastly, we repeated these simulations using two additional TCRs. When using the F24 TCR interacting with an HIV Gag-derived peptide presented on HLA-DR11 (31), we found the same negative correlation with a dimensionless slope of -0.041 (Figure S5A-B). When using the TK3 TCR interacting with an EBV peptide presented on HLA-B*35:01 (32), we found a more modest negative slope of -0.024 (Figure S5C,D).

Taken together, molecular simulations support a negative correlation between x_β_ and *k*_off_ with the quantitative slope varying across different TCR/pMHC complexes. As with our experimental data, this result implies that pulling forces will impair antigen discrimination by reducing fold-differences in off-rates because higher-affinity interactions will be more sensitive to force compared to lower-affinity interactions.

### The OT-I TCR binds OVA pMHC with a low-affinity and an exceptionally fast off-rate

Previous studies used the bioforce probe to report a catch bond for the OT-I TCR binding its agonist OVA pMHC ligand (10, 14). We confirmed this in our laminar flow chamber assay (Figure S6A-B). Since the negative correlation between x_β_ and *k*_off_ predicts that lower-affinity interactions with fast off-rates are likely to be resistant to or even benefit from force, we considered whether the OT-I/OVA interaction may have an unusually low affinity and fast off-rate. While early studies reported high affinities and slow off-rates at 37°C for the OT-I/OVA interaction (*k*_off_ *∼* 0.02 s^*−*1^) (33), more recent measurements reported lower affinities and faster off-rates, even though they were performed at 25°C (14, 34). This discrepancy prompted us to repeat these measurements using an SPR protocol optimised for low affinities at 37°C (4).

We injected purified OT-I over a surface with low levels of purified OVA pMHC (Figure S6C-F) and obtained an affinity of K_D_ = 34 *µ*M (Figure S6C-E), which is unusually weak for TCR interactions with an MHC-I restricted agonist (35). The dissociation phase produced very fast off-rates that were at the SPR instrument limit (Figure S6F) and therefore, we repeated the measurements using a different instrument based on Grating-Coupled Interferometry (GCI) (Figure S6G,H). Both GCI and SPR produced comparably fast off-rates of 4.3 and 5.7 s^*−*1^, respectively (Figure S6I). Using the measured off-rate and affinity we calculated that the on-rate for this interaction was 0.13 *µ*M^*−*1^s^*−*1^ (Figure S6J), which is within the range obtained with other TCRs (35).

Our analysis shows that the OT-I/OVA interaction, which forms a catch bond, has an unusually fast off-rate for an agonist TCR/pMHC interaction (35), 13-17-fold faster than the 1G4/9V interaction (1). This is consistent with our findings that TCR/pMHC interactions with fast off-rates (lower-affinity) are more likely to be resistant to forces (weak slip bonds) or benefit from them (catch bonds).

### Enhanced antigen discrimination by adhesion interactions can be explained by ‘force-shielding’ TCR/pMHC interactions

It is well established that engagement of the T cell adhesion receptors CD2 (which binds CD58) and LFA-1 (which binds ICAM-1) improve the sensitivity of T cells to antigens (9, 18, 23, 36). Recently, we reported that this improvement in sensitivity was progressively lost as the antigen affinity was lowered and consequently, engagement of these receptors increased the ability of T cells to discriminate antigens (4) but the mechanism remains unclear.

To address this, we first modified the standard kinetic proofreading model by including the impact of force (Figure 4A). In the standard model, the duration of pMHC binding to the TCR, which critically determines whether binding is translated to a productive TCR signal, is determined by the solution off-rate. Here, we were able to replace the solution off-rate with the predicted membrane off-rate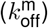using the force sensitivity relationship that we had measured for the 1G4 TCR (Figure 2F, *x*_*β*_ = *−*0.031 log(*K*_*D*_*/*762)). We found that reducing molecular forces from 100 to 10 pN increased differences in signalling TCRs between three test ligands (Figure 4B). This effect is illustrated when the concentration of ligand (*P*) required to produce a threshold level of TCR signal is plotted over the zero-force *k*_off_ (Figure 4C). It is evident that *P* increases faster for lower values of *k*_off_ compared to higher values under an applied force. Consequently, the model predicts that increasing force on the TCR/pMHC interaction decreases antigen sensitivity (Figure 4D) and reduces antigen discrimination (Figure 4E).

**Figure 4.**
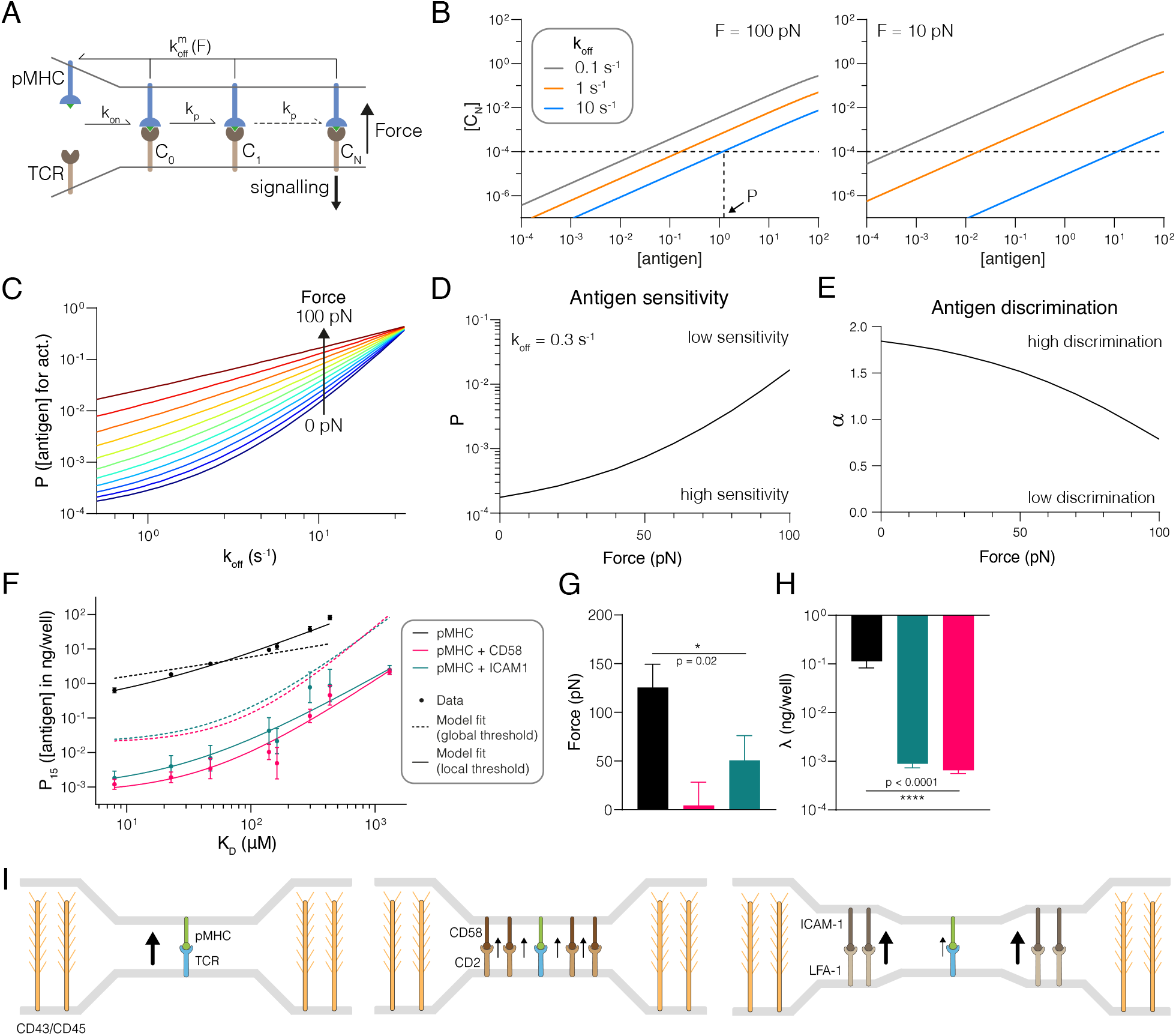
The ability of adhesion receptors to enhance T cell antigen discrimination and sensitivity can be explained by them shielding the TCR/pMHC interaction from mechanical force. Schematic of the kinetic proofreading model modified to allow molecular forces to impact the membrane off-rate 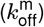. **(B)** The concentration of productively signalling TCRs (*C*_*N*_) over the antigen concentration for antigens with the indicated zero-force off-rates (*k*_off_). Shown is the effect of forces at 100 pN (left) and 10 pN (right). The horizontal dashed line is the threshold value of *C*_*N*_ required for activation in the model. The vertical dashed line shows the antigen concentration (*P*) required to elicit activation by the antigen with the largest value of *k*_off_. **(C)** The antigen concentration required to elicit activation (*P*) over *k*_off_ for the indicated molecular forces on the TCR/pMHC interaction. **(D)** The value of *P* over the applied force for Ligand A defined in panel (C). **(E)** The discrimination power (*α*) over the applied force. Discrimination power is defined as the ratio of the logarithm of the fold-change in *P* over the fold-change in *k*_off_ for Ligand A and B defined in panel (C). **(F)** Experimental value of *P* (symbols) over antigen affinity fitted by the kinetic proofreading model with force (dashed and solid lines). The value of *P* is defined as the concentration of pMHC required for 15% upregulation of CD69 above baseline and is shown as the mean with SEM from 4 independent experiments. All experimental data is taken from Fig. 4 in (4). **(G)** The fitted value of the applied force. **(H)** The fitted value of *λ*. An F-test is used to produce a p-value for the null hypothesis that the applied force or *λ* is the same across the three conditions. All calculations (B-E) and data fitting (F-H) used *N* = 2.46 and *k*_*p*_ = 2.15 s*−*1 (average of CD58 and ICAM-1 conditions) taken from Fig. 5 in (4). **(I)** Proposed mechanisms by which adhesion receptors can reduce pulling forces on TCR/pMHC interactions (black arrow). Molecular forces on individual TCR/pMHC interactions can be produced by multiple mechanisms, including a mismatch in size between the short TCR/pMHC interaction (*∼* 14 nm) and larger surface molecules, such as CD43 and CD45 that extend 21-45 nm (left panel). These forces may be reduced by force-sharing with the CD2/CD58 adhesion molecules that span the same distance and can therefore closely colocalise (middle panel). Alternatively, the larger LFA-1/ICAM-1 adhesion interaction may reduce forces by acting as a diffusional barriers to maintain CD43 and CD45 further away from TCR/pMHC interactions, and tether and stabilize cellular processes such as microvilli, thereby reducing the forces imposed on all interactions within an area of close contact surrounded by LFA-1/ICAM interactions (right panel)

To understand how adhesion receptors impact the force on the TCR/pMHC interaction, we next fitted the modified model to data we previously generated (4). Briefly, primary human CD8^+^ T cells expressing the 1G4 TCR were stimulated with eight antigens either alone or in combination with CD58 or ICAM-1, and the concentration of antigen required to elicit activation (*P*) was plotted against the solution affinity (Figure 4F). We fixed the number (*N*) and rate () of each step to those we previously identified (4) to obtain a mechanical proofreading model with only two fitting parameters: the force applied on the TCR/pMHC interaction (*F*) and a quantity that is proportional to the threshold concentration of productively signalling TCRs (*λ*). Intuitively, lower values of *λ* mean that a lower TCR signal is sufficient to activate a cellular response.

We first asked whether differences in force are sufficient to explain the changes in *P* for all antigens. To do this we fitted *F* while constraining *λ* to a single ‘global’ value (Figure 4F, dashed line). We found that, while the fit qualitatively reproduced the data, in that ligation of CD2 or LFA-1 increased both antigen sensitivity and discrimination, it failed to quantitatively fit the data. Inspection of the fits suggests that force shielding by CD2 and LFA-1 (dashed lines) can fully account for their ability to enhance antigen discrimination (changes in slope) but only partially account for their ability to enhance antigen sensitivity (vertical shifts/changes in potency). When we also allowed *λ* to vary, we observed an excellent fit (Figure 4F, solid line) with reduced TCR/pMHC forces (Figure 4G) and reduced TCR signalling thresholds (Figure 4H) upon ligation of adhesion receptors.

In summary, these results suggest that force-shielding by CD2 and LFA-1 can fully account for their ability to improve antigen discrimination and partially account for their ability to enhance sensitivity, suggesting that sensitivity is further enhanced by processes, presumably signalling by CD2 and LFA-1, that amplify TCR signals.

## Discussion

To understand how molecular forces impact antigen discrimination, we used Bell’s model to show that discrimination is impaired under force when the TCR/pMHC sensitivity to force (x_β_) is negatively correlated with *k*_off_. Using cell-free experiments, we confirmed this prediction showing that higher-affinity interactions are more susceptible to force (slip bonds with a higher x_β_) than lower-affinity interactions that were also more likely to form catch bonds (x_β_*<*0), and consistent with this, we found that the best-characterized catch bond (OT-I/OVA) has an unusually low affinity and the fastest *k*_off_ yet reported for an agonist TCR/pMHC interaction. Molecular dynamics simulations of these and other TCR/pMHC interactions confirmed this correlation, suggesting that it may be a general feature of TCR/pMHC interactions.

Our findings contrast with previous reports suggesting that force on the TCR increases antigen discrimination (10, 25). These reports found that high-affinity agonist pMHC formed catch bonds whereas low-affinity non-agonists pMHC formed slip bonds. Importantly, these studies mostly relied on measurements using the bioforce probe, where the TCR is on T cell so that repeated engagement by agonist pMHC will trigger the T cell. Furthermore, increasing the force on the TCR enhances TCR triggering (9, 37, 38), which induces cellular responses likely to enhance TCR/pMHC binding, such as coreceptor recruitment, TCR aggregation, and changes in the cytoskeleton or membrane. For example, initial TCR binding has been shown to enhance subsequent binding events in this system (39), and CD8 recruitment induced by TCR triggering has been shown to stabilize the TCR/pMHC interaction (40). Since this can confound measurements of the effect of force on *k*_off_ it is important to verify measurements in a cell-free system. Indeed, when the same TCR/pMHC interactions have been studied in a cell-free system their sensitivity to force was increased (14). This includes the 1G4 TCR, which is reported to have a catch bond using the bioforce probe (12) but a slip bond in a cell free system [this study and (26)]. These results suggest that the greater resistance of TCR/pMHC interactions to force reported using intact T cells is a consequence of TCR triggering. This would account for the fact that, in these systems, agonist pMHC ligands exhibit catch bonds whereas non-agonist pMHC form slip bonds.

In addition to a constant force modifying T cell antigen discrimination through variation in force sensitivity (x_β_), it has been proposed that *changes* in force during TCR/pMHC engagement can improve T cell antigen discrimination. Allard et al showed that a decreasing time-dependent force would increase antigen discrimination by preferentially rupturing antigens with fast *k*_off_ (19). Such a situation could arise if large molecules are initially trapped and compressed within and then diffuse out of the close contact areas where TCR engages pMHC. It has also been argued that ramping up the force on a TCR/pMHC interaction could improve antigen discrimination by narrowing the probability distribution of bond-lifetimes (41). This could occur if the TCR and pMHC were moved apart from each other at a constant velocity as a result, for example, of the movement of the cells or their processes. Our findings do not rule out a role for either of these mechanisms in modulating the discriminatory power of the TCR.

Why might TCR/pMHC interactions with a lower affinity/faster *k*_off_ be more resistant to force than higher affinity interactions? One intuitive explanation is that the contact interface will be less intimate with low versus high affinity interactions. It follows that the notional distance between the energy minimum and the energy barrier along the dissociation pathway would be shorter. Thus x_β_, which has units of length and can be thought of as equivalent to this distance, will be shorter for low versus high affinity interactions. Assuming that the height of the energy barrier decreases less than this distance for lower-affinity interactions, shortening will make the slope up to the energy barrier steeper and, since this slope can be considered to be equivalent to force, this implies greater force is needed to reach the barrier. While the unbinding pathway for protein/proteins interactions is likely to be complex and varying, it seems reasonable that mutations within a given TCR/pMHC complex that increase or decrease the affinity would increase or decrease respectively, the intimacy of the contact interface and thus x_β_. In contrast, given the large structural diversity between TCR/pMHC binding interfaces (42), and the fact that TCRs do not encounter foreign agonist pMHC during their development, it is difficult to envisage a molecular explanation for high-affinity interactions all forming catch bonds. Instead, our data suggests that catch bonds formed by agonists, such as OT-I/OVA, will have unusually fast *k*_off_ so that they can only function as agonists by forming a catch bond that sufficiently slows their 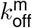 to induce TCR signals. Consistent with this, a recent mutagenesis screen for TCR/pMHC interactions that were activatory despite having low affinities identified several apparent catch bonds, as measured by the bioforce probe (13). The structural explanation for the x_β_-*k*_off_ relationship described above cannot easily explain catch bonds and why they may be more common with low-affinity interactions. One possible explanation is that lower affinity TCR/pMHC complexes possess more conformational flexibility within the binding interface, increasing the likelihood of conformational adjustments leading to new interactions during unbinding under force.

We have previously shown that the adhesion molecules CD2 and LFA-1 improve T cell antigen discrimination. Here, we have found that the kinetic proof-reading mechanism, when modified to include the effects of force, can explain the effect of CD2 and LFA-1 on antigen discrimination by their ability to shield the TCR/pMHC interaction from forces. This is readily explained in the case of CD2 because CD2/CD58 interactions span the same distance as the TCR/pMHC interaction. This size compatibility enables them to closely colocalise with individual TCR/pMHC interactions within close contacts and hence share forces (Figure 4I, left and middle panels). Indeed, modelling confirms that receptor/ligand interactions can share forces when co-localised on the nanometer scale (43, 44). Given that the LFA-1/ICAM-1 interaction spans a greater distance than the TCR/pMHC interaction (45), and does not intimately colocalize with it within the immunological synapse (46), it cannot reduce TCR/pMHC forces by the same CD2/CD58 force shielding mechanism. Instead, we suggest that LFA-1/ICAM-1 interactions reduce force on a larger scale, by acting as a diffusional barrier to exclude large glycocalyx molecules such as CD43 and CD45 (19**?**), and through their ability to reduce T cell mobility and/or stabilise lamelopodia and microvilli processes (e.g. Figure 4I, right panel). In support of this, LFA-1/ICAM-1 interactions have been shown to form ‘micro-adhesion’ rings surrounding areas of TCR/pMHC engagement (46). Furthermore, LFA-1/ICAM-1 interactions experience substantial forces (*>*56 pN) at T cell interfaces, and increasing their ability to withstand forces enhances antigen discrimination (7). Direct measurements of the effect of engaging adhesion receptors on the force experienced by the TCR at T cell/APC interfaces are needed to test these predictions.

In conclusion, we have shown that force sensitivity increases with affinity, which implies that force on the TCR/pMHC interaction impairs antigen discrimination. The fact an antibody/antigen interaction shows the same correlation suggests that this may be general feature of antigen receptors. Importantly, B cell antigen recognition also occurs at cellular interfaces where mechanical forces are likely to play an important role (47). Given the critical role of T and B cell antigen discrimination, we suggest that force shielding is functionally important as it provides an environment where antigen receptors experience a consistent level of force. The fact that CD2 and LFA-1 are expressed on all T cells, and their ligands CD58 and ICAM-1 are ubiquitously expressed, makes them suitable for such roles.

## Acknowledgements

We thank Edward A. FitzGerald for technical assistance with GCI and Jun Allard for discussion on adhesion receptor mechanics. We thank Bernhard Knapp, Dheeraj Prakaash, and Antreas Kalli for initial discussions on molecular dynamics simulations.

## Open access

This research was funded in whole, or in part, by the Wellcome Trust [207537/Z/17/Z]. For the purpose of Open Access, the author has applied a CC BY public copyright licence to any Author Accepted Manuscript version arising from this submission.

## Methods

### Laminar flow assay

Glass slides were rinsed twice in absolute ethanol then washed in a “piranha” solution composed of 70 % H_2_SO_4_, 15 % water and 15 % H_2_O_2_ for 10 minutes, then rinsed with 5 liters of deionized water. Glass slides were coated with poly-L-lysine (150–300 kDa, Sigma Aldrich, France) at 100 *µ*g/ml in a 0.01M phosphate solution, pH 7.4 for 10 minutes, then rinsed in PBS, then incubated with glutaraldehyde (2.5 % in pH 9.5 0.1 M borate solution, Sigma Aldrich) for 10 minutes, then rinsed in PBS, then incubated in a 100 *µ*g/ml biotinylated BSA solution in PBS (Sigma Aldrich), for 30 minutes, then rinsed in PBS, then incubated in a 10 *µ*g/ml streptavidin solution in PBS (Sigma Aldrich), then rinsed in PBS. Glass slides were then mounted in a home-made multi-channel thermo-regulated flow chamber device, forming nine independent chambers of 12 mm length and 2×0.250 mm^2^ section. Each chamber was then filled with a biotinylated pMHC solution in PBS with 0.1 % BSA at chosen concentration, using cascade dilution to deposit in our 9-chambers apparatus 8 different amounts of pMHC in each chamber plus one chamber as a negative control. Microspheres (Dynal M-450 tosly-activated, ThermoFisher, France) were rinsed three time in a 0.1 M borate solution, then incubated in a 200 *µ*g/ml solution of anti-6xHis-tag antibody (Thermo Fisher) overnight under constant agitation. Microspheres were rinsed three time in PBS/0.1 % BSA then incubated prior to each experiment in a 200 *µ*g/ml solution of 6xHis-tag TCR.

Flow chamber experiments were performed using our automaton based on a Arduino Mega 2560 card (Arduino, Italy). The device forming nine independent chambers on a common glass slide was thermo-regulated at 37°C by water circulation, set on an inverted microscope with a 10x lens (Leica, Germany) with a digital CCD camera (UEye, IDS Germany), and chamber entry was connected to the piping. For each independent chamber, the automaton performed cycles, each cycle being an experiment for a given shear flow. The automaton repeated cycles of microspheres agitation, microspheres injection in the chamber, launch of movie recording at 50 frames per seconds with M-JPEG on-the-fly compression, and flow at a given shear for 90 seconds. Shear value was automatically modified for each new cycle until all chosen shear conditions had been recorded. The chamber was manually disconnected and next chamber was connected, then automaton was re-launched. Raw data are in the form of movies of microspheres motion. A software suite written in Java as ImageJ plug-ins retrieve microspheres trajectories, then detects microspheres arrest events using a velocity threshold and record arrests number, arrests duration and distance traveled by sedimented microspheres.

Linear binding densities are calculated as the ratio of the number of arrests on the distance traveled by microspheres sedimented on the chamber surface. Specific binding densities are calculated by subtracting control linear binding density from assay linear binding density for a given shear condition. Duration of arrests were pooled for experiments sharing identical TCR and pMHC molecules, identical amount of pMHC on the surface and identical shear rate, to built survival curves of the arrests. Data points that did not have more events than the corresponding control experiment were excluded. Each TCR-pMHC bond was measured in 8 to 12 independent experiments. Single molecular bond observation was assessed using the usual flow chamber arguments: in an interval of deposited amounts of pMHC, linear binding density was increasing linearly from negative control value with the mount of deposited ligand; survival curves of bonds would not change in the same range of amounts of deposited pMHC, showing observation of similar binding events. For this binding density analysis, specific survival curves were calculated by subtracting, for each time step, the corresponding survival fraction of non-specific arrests measured in control experiments.

### Data analysis

Pre-processed data from the laminar flow chamber assay was analyzed using a custom Python pipeline (Python 3.8.3 AMD64, lmfit 1.0.1, matplotlib 3.2.2) and GraphPad Prism 9.3.0 (GraphPad Software). Experiments with less than 20 events in the first second were excluded. For each time point with at least one recorded event, the survival fraction was calculated. To calculate off-rates, we determined the survival fraction at 1 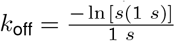 In the rare cases where there was no event at exactly 1 s, we used a subsequent event. We fitted Bell’s model (24) to each 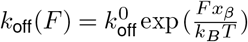, where F is the force in pN, 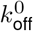the zero-force off-rate (s^-1^; fitted parameter), *x*_*β*_ the force sensitivity (nm; fitted parameter), *k*_*B*_ the Boltzmann constant 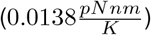, and *T* the temperature (310.15 K).

**Table 1.**
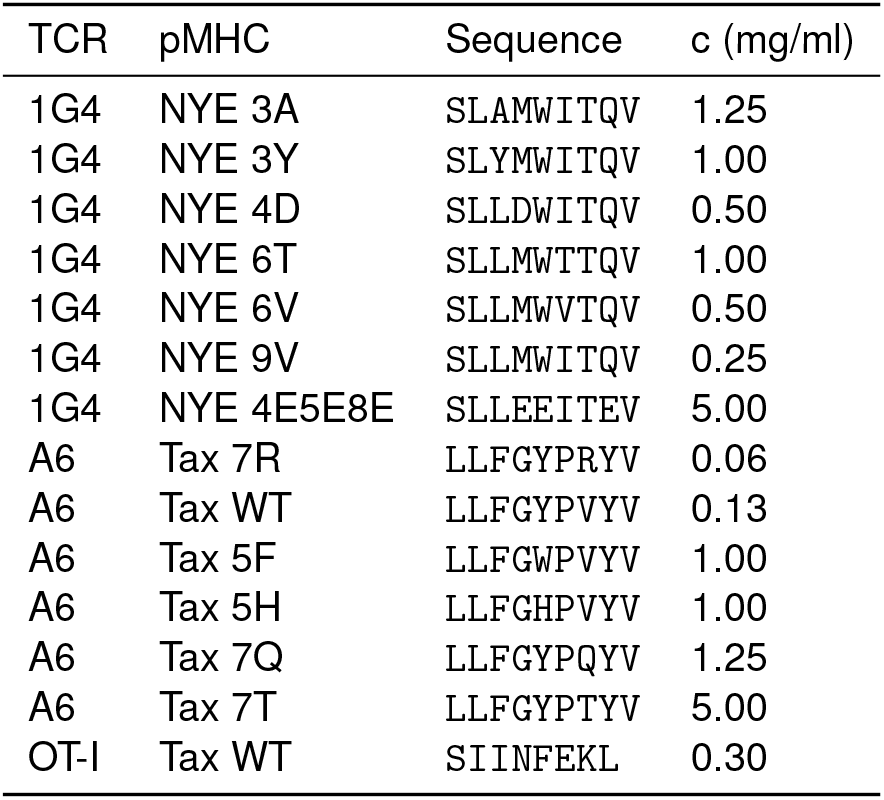
pMHC concentrations used.

### Protein expression and purification

Soluble OT-I TCR construct consisted of the murine variable OT-I domain and the human constant domain truncated above the trans-membrane domain with an artificial interchain disulphide. Soluble 1G4 TCR (no artifical disulphide) and A6 TCR (with an artificial disulphide) were similarly truncated above the trans-membrane domain (TRAC residue 93, TRBC2 residue 129). All TCRs contained a 6xHis-tag on one chain to allow immobilization on beads for force experiments. TCR α and β chains were expressed in BL21 (DE3)-RIPL *Escherichia coli* cells (Agilent Technologies) following induction with 0.15 mM IPTG. Inclusion bodies were isolated by disrupting cells with BugBuster (Merck) according to the manufacturer’s instructions. Proteins were stored at -80°C until use. TCRs were refolded by adding 15 mg (OT-I) or 30 mg (1G4 or A6) of each chain dropwise in 1 L cold refolding buffer (0.15 M Tris-HCl pH 8.0, 3 M urea, 0.2 M Arg-HCl, 0.5 mM EDTA), followed by dialysis for 3 days in 10 L dialysis buffer (10 mM Tris-HCl pH 8.5), with a buffer change after day 1. After dialysis, the protein was filtered and purified using anion-exchange chromatography (HiTrap Q column [GE Healthcare]) with a NaCl gradient, followed by concentration and purification by size exclusion chromatography (Superdex 200 column [GE Healthcare]) in HBS-EP (0.01 M HEPES pH 7.4, 0.15 M NaCl, 3 mM EDTA, 0.005 % v/v Tween20). TCR purity was checked by SDS-PAGE and concentrations were measured with a Nanodrop spectrophotometer (Thermo Fisher). Purified TCR were stored at 4°C and used for SPR and GCI measurements withing 24 h (OT-I) or 1 month (1G4 and A6) after purification to avoid aggregation.

Soluble class I pMHCs bound to OVA peptide (OVA_257–264_; SIINFEKL) or A2 variant (SAINFEKL) were refolded and biotinylated by the NIH protein facility. We used mouse H-2Kb heavy chain and human beta-2 microglobulin, biotinylated on the C terminal of the heavy chain. pMHCs were aliquoted and stored at -80°C until use.

NYE (NYE-ESO_157–165_) and Tax (HTLV-1 Tax_11–19_) class I pMHCs were refolded in-house. The heteroclitic 9V variant was used as index peptide for the 1G4 TCR, rather than the wild-type 9C, due to its improved stability on MHC (48). Soluble human HLA-A*0201 heavy chain (UniProt residues 25–298) with a C-terminal AviTag/BirA recognition sequence and human beta-2 microgolublin were expressed in *Escherichia coli* and isolated from inclusion bodies. Trimer was refolded by consecutively adding peptide, *β*2M and heavy chain into refolding buffer and incubating for 2–3 days at 4°C. Protein was filtered, concentrated using centrifugal filters, biotinylated (BirA biotin-protein ligase bulk reaction kit [Avidity, USA]) and purified by size exclusion chromatography (Superdex 75 column [GE Healthcare]) in HBS-EP. Purified protein was aliquoted and stored at -80°C until use.

Soluble extracellular domain (ECD) of human CD58 (UniProt residues 29-–204 + AviTAG + 6xHis) and human CD86 (UniProt residues 24–238 + AviTAG + 6xHis) were produced in Freestyle 293F suspension cells (Thermo Fisher) according to the manufacturer’s instructions. Ligands were biotinylated by co-transfection (1:10) of a secreted BirA-encoding plasmid (pTT3-BirA-FLAG) and adding 100 *µ*M D-biotin to the medium, as described before (49). All supernatants were 0.45 *µ*m filtered and 100 *µ*m PMSF was added. Proteins were purified using standard Ni-NTA agarose columns. Proteins were further purified by size exclusion chromatography (Superdex 75 or 200 column [GE Healthcare]) in HBS-EP; purified proteins were aliquoted and stored at –80°C until use.

### Surface plasmon resonance

TCR–pMHC interactions were analyzed on a Biacore T200 instrument (GE Healthcare Life Sciences) at 37°C and a flow rate of 10 *µ*l/min. Running buffer was HBS-EP. Streptavidin was coupled to CM5 sensor chips using an amino coupling kit (GE Healthcare Life Sciences) to near saturation, typically 10000–12000 response units (RU). Biotinylated pMHCs (47 kDa) were injected into the experimental flow cells (FCs) for different lengths of time to produce desired immobilization levels (typically 500–1500 RU), which were matched as closely as feasible in each chip. Usually, FC1 was as a reference for FC2–FC4. Biotinylated CD58 ECD (24 kDa + *∼*25 kDa glycosylation) was immobilized in FC1 at a level matching those of pMHCs. In some experiments, another FC was used as a reference. Excess streptavidin was blocked with two 40 s injections of 250 *µ*M biotin (Avidity). Before injections of soluble 1G4 or A6 *αβ*TCR (51 kDa), the chip surface was conditioned with 8 injections of the running buffer. Dilution series of TCRs were injected simultaneously in all FCs; the duration of injections (30–70 s) was the same for conditioning and TCR injections. After every 2–3 TCR injections, buffer was injected to generate data for double referencing. After the final TCR injection and an additional buffer injection, W6/32 antibody (10 *µ*g/ml; Biolegend; Lot: B233942) was injected for 10 min.

TCR Steady-state binding was measured >10 s post-injection. In addition to subtracting the signal from the reference FC with immobilized CD58 (single referencing), all TCR binding data was double referenced vs. the average of the closest buffer injections before and after TCR injection. This allows to exclude small differences in signal between flow cells (e.g. drifts). TCR binding versus TCR concentration was fitted with the following model: *B* = *B*_*max*_*∗* [*TCR*]*/*(*K*_*D*_ + [*TCR*]), where *B* is the response/binding, B_max_ the maximal binding (this parameter is either kept free or is fixed with the W6/32 derived B_max_), and [*TCR*] the injected TCR concentration. Maximal W6/32 binding (R_max_) was used to generate the empirical standard curve and to infer the B_max_ of TCRs from the standard curve. R_max_ was derived by fitting the W6/32 binding data after double referencing with the following, empirically chosen, model: *R* = *R*_*max*_ *∗ t/*(*K*_*t*_ + *t*), where *t* is time (s), *R* the sensogram response after single referencing, and *K*_*t*_ a nuisance parameter. The empirical standard curve only contained data where the ratio of the highest concentration of TCR to the fitted K_D_ value (obtained using the standard method with B_max_ fitted) was 2.5 or more. This threshold ensured that the binding response curves saturated so that only accurate measurements of B_max_ were included. All interactions were fit using both the fitted and constrained B_max_ method. For constrained K_D_ above 20 *µ*M we reported the constrained K_D_, otherwise we use the B_max_ fitted K_D_. SPR data was analyzed using GraphPad Prism 8 and 9 (GraphPad software) or using a custom Python script (Python v3.7 and lmfit v0.9.13).

### Surface plasmon resonance and grating-coupled interferometry experiments for OT-I TCR

Binding properties of OT-I TCR interaction with OVA were measured by SPR on a Biacore S200 and T200 (GE Healthcare Life Sciences) using a CM5 sensor chip and by GCI on the WAVEsystem (Creoptix) with a 4PCP sensor chip. HBS-EP was used as running buffer and all measurements were performed at 37°C. For protein immobilization, the sensor chip was saturated with streptavidin using an amine coupling kit (GE Healthcare Life Sciences). Biotinylated pMHCs were immobilize at various levels (100–300 RU for kinetics, 1000–2000 RU for affinity measurements). CD86 with matching immobilization levels were use as reference protein. Excess streptavidin was blocked with two 40 s injections of 500 *µ*M biotin (Avidity) and the sensor was conditioned with at least 8 injections of running buffer. TCR concentrations used varied between 50–150 *µ*M.

Binding affinities were measured by equilibrium binding on a T200 Biacore instrument. TCR was injected at increasing concentrations at 30 *µ*l/min with flow path 1-2-3-4. Buffer was injected after every 2–3 TCR injections. K_D_ values were obtained by fitting a 1:1 Langmuir binding model (*RU*_eq_ = *RU*_max_ *×* [*TCR*]*/*(*K*_*D*_ + [*TCR*])) to double-referenced equilibrium RU values.

For kinetic measurements by SPR we used a Biacore S200. Different TCR concentrations were injected at a flow rate of 30 *µ*l/min. To minimize diffusion artifacts, TCR was injected separately in flow path 1-2 and 3-4. We obtained *k*_off_ by fitting a mono-exponential to double referenced dissociation curves.

For kinetic measurements by GCI we used a Creoptix waveRAPID. A single TCR concentration was injected multiple times using different length pulses for a total duration of 5 s, followed by a 50 s dissociation phase. To calibrate how the analyte (TCR) concentration changes over time during pulse injection, 0.5 % DMSO is injected. Flow rate was set to of 100 *µ*l/min per flow cell. We obtained *k*_off_ by fitting a mono-exponential to double referenced dissociation curves.

### Coarse-grained molecular dynamics simulations

We have used an implicit-solvent coarse-grained model in which amino acid residues are represented by single beads (29). The beads are tethered together into chains by harmonic springs. Non-local interactions between the beads are introduced on the basis of the PDB structure of the protein complex under study. Specifically, interactions between the beads that form contacts in the PDB structure of the protein complex are described by the Lennard-Jones potential. Interactions between the beads that do not form contacts in the PDB structure are purely repulsive and short-ranged. Disulphide bonds are captured by harmonic springs between the specified Cys beads. Importantly, the amino acid contacts in PDB structures are identified using an overlap criterion applied to the coordinates of all heavy atoms in the structures. The pairs of amino acid residues that are very close sequentially, *i*.*e*. (i,i+1) and (i,i+2), are excluded from the set of contacts in the PDB structure. A detailed description of the coarse-grained model is given in (29).

As input to our simulations we used the PDB structures of the 1G4 and A6 TCR-pMHC complexes with the PDB codes 2BNQ and 3QFJ, respectively. We considered five constructs of pMHC in complex with the 1G4 TCR, where (i) all contacts between the peptide and the TCR identified in the PDB structure (PDB code: 2BNQ) were included in the coarse-grained model, (ii) contacts between the peptide residues 4 to 8 and the TCR identified in the PDB structure were excluded from the coarse-grained model, (iii) only contacts between the peptide and the TCRα identified in the PDB structure were taken into account in the coarse-grained model, (iv) only contacts between the peptide and the TCRβ identified in the PDB structure were taken into account in the coarse-grained model, and (v) no contacts between the peptide and the TCR identified in the PDB structure were included in the coarse-grained model. We also studied four constructs of pMHC in complex with the A6 TCR, where (i) all contacts between the peptide and the TCR identified in the PDB structure (PDB code: 3QFJ) were included in the coarse-grained model, (ii) only contacts between the peptide and the TCRα identified in the PDB structure were taken into account in the coarse-grained model, (iii) only contacts between the peptide and the TCRβ identified in the PDB structure were taken into account in the coarse-grained model, and (iv) no contacts between the peptide and the TCR identified in the PDB structure were included in the coarse-grained model.

We performed molecular dynamics simulations of the coarse-grained model using the Langevin thermostat. All of the simulations were started from native states corresponding to the PDB structures. Stretching of the TCR-pMHC complexes was implemented by attaching harmonic springs to two beads: the first one to the C-terminus of the TCRβ and the second one to the C-terminus of the MHC. The first of the springs was fixed in space and the second one was moved with a constant speed v. We performed the simulations with v ranging from 0.0005 to 0.005 Å/τ, where the time unit τ is estimated to be of the order of 1 ns. To prevent unfolding of the individual chains within the TCR-MHC complexes, we replaced the intra-molecular contacts by harmonic springs. The inter-molecular contacts, however, were still described by the Lennard-Jones potentials, as introduced in (29). This modification resulted in *F* (*t*) traces with single peaks (Figure 3A–B) that were identified to coincide with TCR-pMHC dissociation events.

We monitored the response force *F* acting on the pulling spring (Figure 3A) and determined the average force *F*_max_ at which the TCR-pMHC complexes break apart. The average was obtained from twenty independent trajectories (Figure 3B). The dependence of *F*_max_ on *ν* follows the Bell-Evans formula (30):

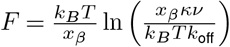

which yields the values of parameters *k*_off_ and x_β_ (Figure 3C). Here, *k*_*B*_ is the Boltzmann constant, *T* is the absolute temperature, and *κ* = 0.12 *k*_*B*_*T/*Å^2^ is the stiffness of the harmonic springs attached to the C-termini of MHC and TCR β. (The product of *κ* and *ν* equals to the loading rate).

### Kinetic proofreading with molecular forces

All calculations in Fig. Figure 4 were performed with the kinetic proofreading model modified to include the membrane off-rate under force. At steady-state, the concentration of signalling TCR in state *N* is calculated to be,

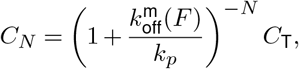

where

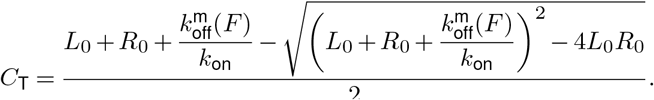

In this model, *L*_0_ and *R*_0_ are the total concentration of pMHC and TCR, respectively, *k*_on_ is the on-rate and 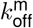 (F) is the membrane off-rate under force (F). We used the empirical relationship determine for the 1G4 TCR for all calculations (Fig. Figure 2F),

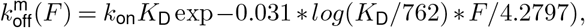

where *k*_on_ was taken to be 0.0447 *µ*M^*−*1^s^*−*1^, which is the average *k*_on_ value for the subset of pMHCs where kinetic parameters were available (see (4) for details). We fixed the value of N to 2.46 and *k*_*p*_ to 2.15 s^*−*1^ to the value we obtained for the plate-data when providing adhesion ligands (Fig. 5 in (4)). Potency (*P*) was defined as the concentration of pMHC (*L*_0_) required to achieve a value of C_*N*_ of 10^*−*4^. Antigen sensitivity was defined as the value of *P* for the highest affinity antigen and antigen discrimination was defined as the logarithm of the fold-change in *P* over the logarithm of the fold-change in *k*_off_ for the highest affinity ligands over the lowest affinity ligands.

To directly fit the model to experimental potency data, we solved for the concentration of pMHC required to obtain a threshold value of *C*_*N*_ (see (4) for complete derivation),

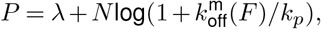

where *λ* is proportional to the threshold value of *C*_*N*_, *N* is the number of proofreading steps, *k*_*p*_ is the rate of each step, and 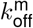 is defined above. In this fit, the only free parameters were *F* and *λ*. We fit the model to the plate-data (pMHC alone, pMHC + CD58, and pMHC + ICAM-1) using Prism (GraphPad) using a different value of *F* for each condition (local force) and either a single value of *λ* (global threshold) or a different value of *λ* for each condition (local threshold).

## Extended Data

**Figure S1.**
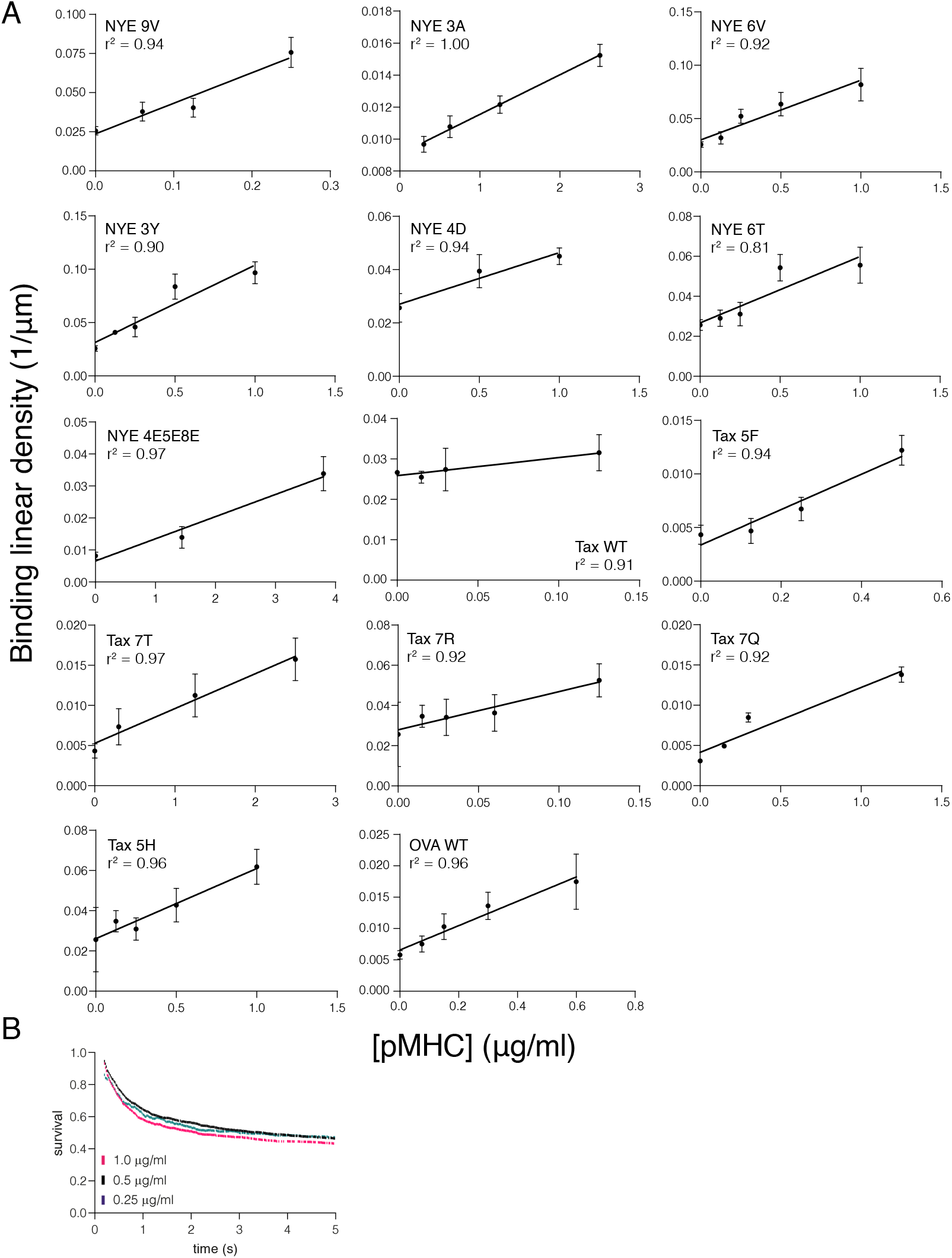
Linear binding density correlation indicates assay operation in single molecule bond regime. **(A)** Linear binding densities for all tested TCR/pMHC combinations. A linear correlation indicates a single-molecule binding regime. **(B)** Example of survival curves of the same TCR/pMHC interaction with different pMHC immobilization levels. Shown is 1G4/NYE 6T.

**Figure S2.**
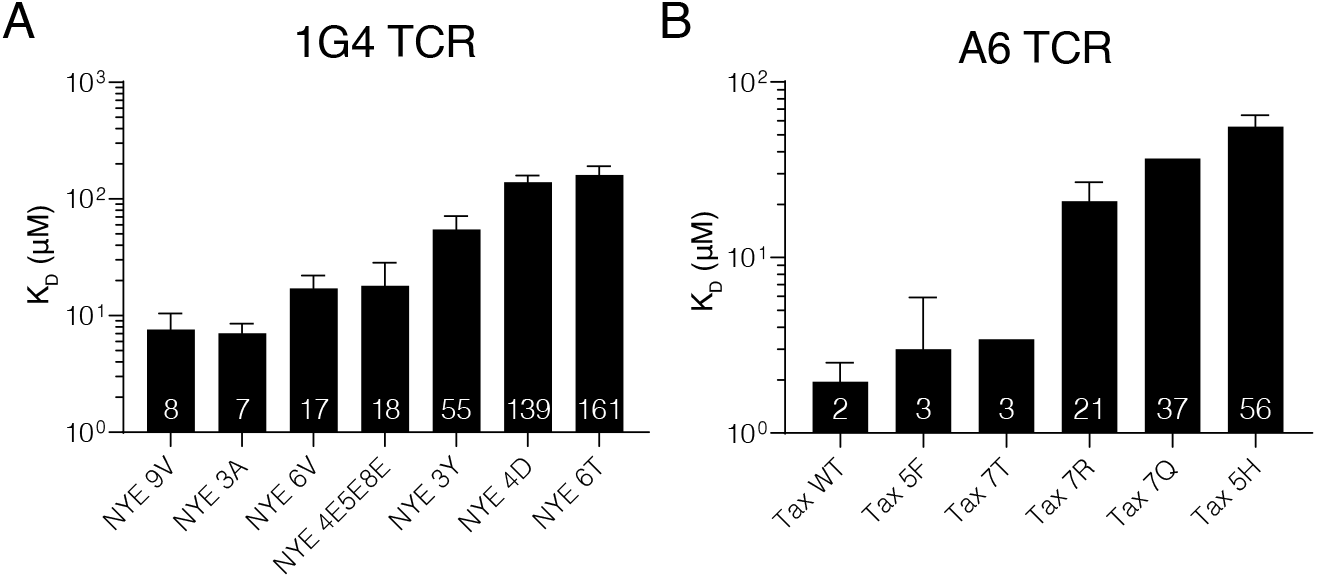
Dissociation constants of TCR/pMHC bond measured by SPR. For **(A)** 1G4 (n=2–10) and **(B)** A6 TCR (n=1–6). Shown are geometric means with SDs. Data is partially reproduced from our previous measurements in (4) with additional repeats and pMHC variants.

**Figure S3.**
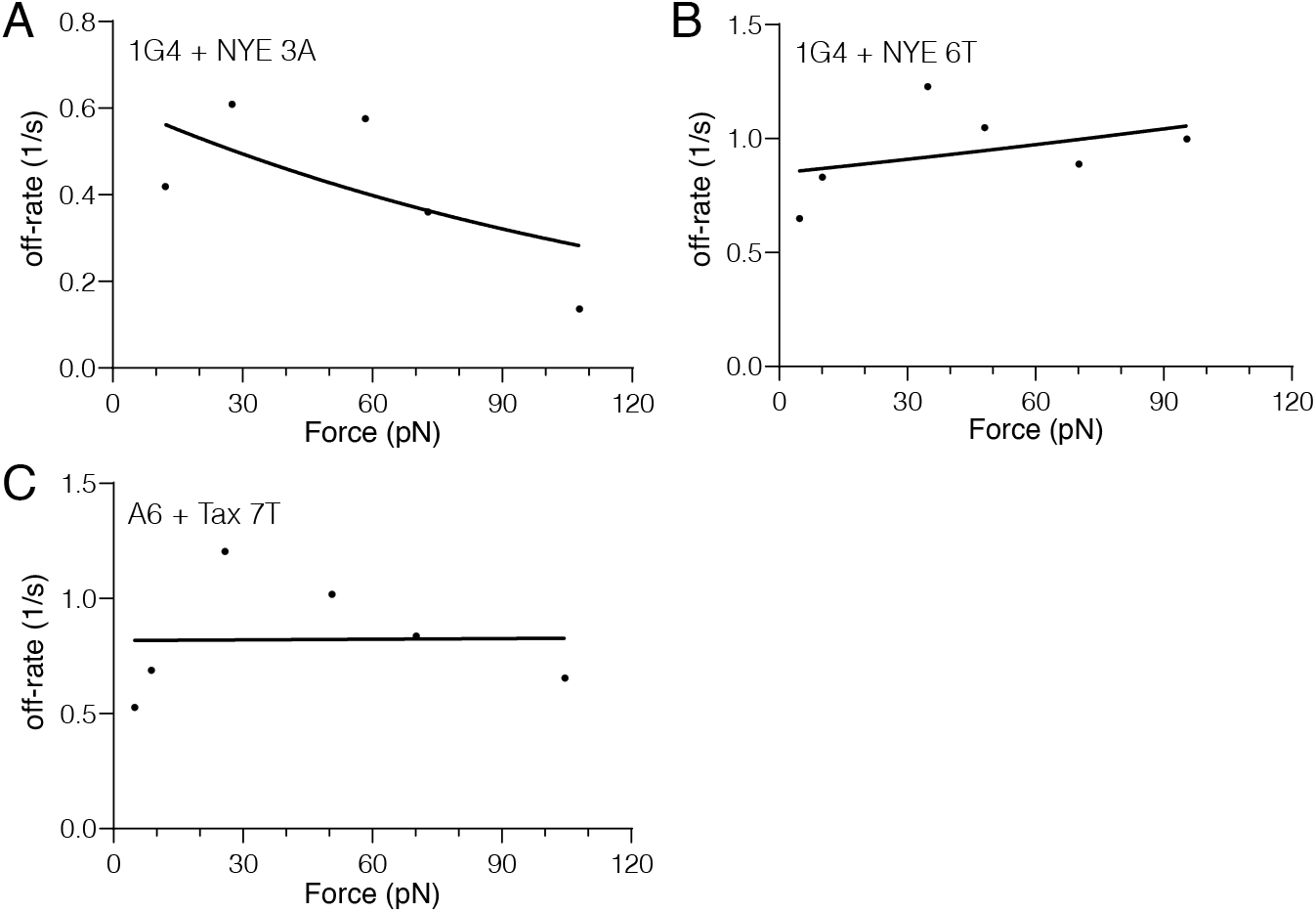
Non-canonical TCR/pMHC bonds. **(A–C)** Data fitted with Bell’s model. Since all of them did not show a classical force-response, as predicted by Bell’s model, we excluded them from any correlation. Notably, they all exhibit slip bond behavior at low forces. The number of independent experiments that were combined to produce the estimated off-rates are: 9 (1G4/3A), 9 (1G4/6T), 11 (A6/7T).

**Figure S4.**
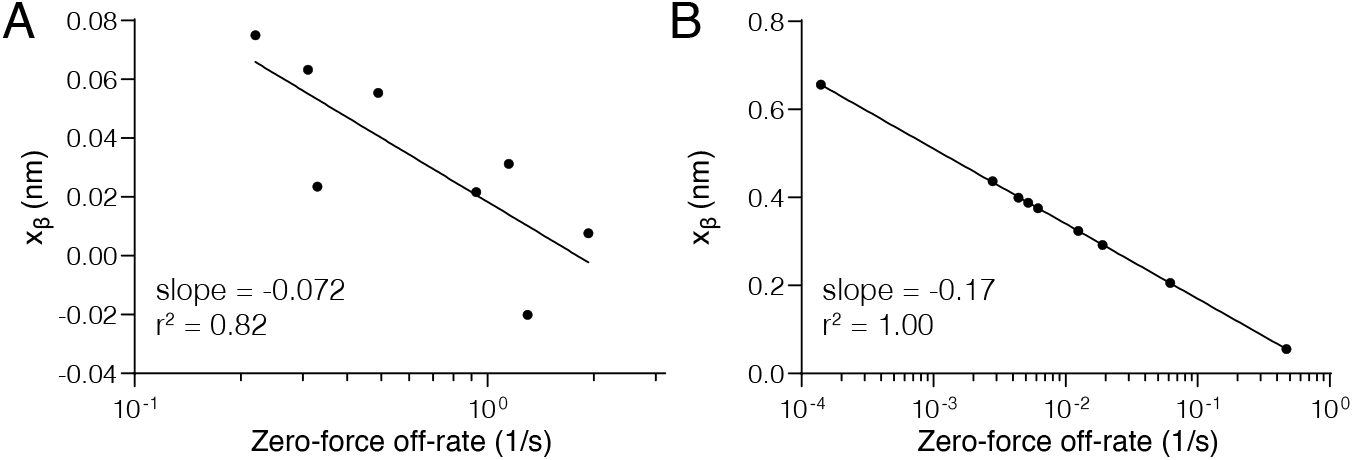
Negative x_β_ k_off_ correlation is reproduced in two other studies. **(A)** We previously generated data at 25°C with a similar correlation using the 1G4 TCR and peptide variants, as well as, MHC molecules with different mutations (26). **(B)** Schwesinger et al. found a negative correlation for different antibodies binding the same antigens when measured by atomic force microscopy (30).

**Figure S5.**
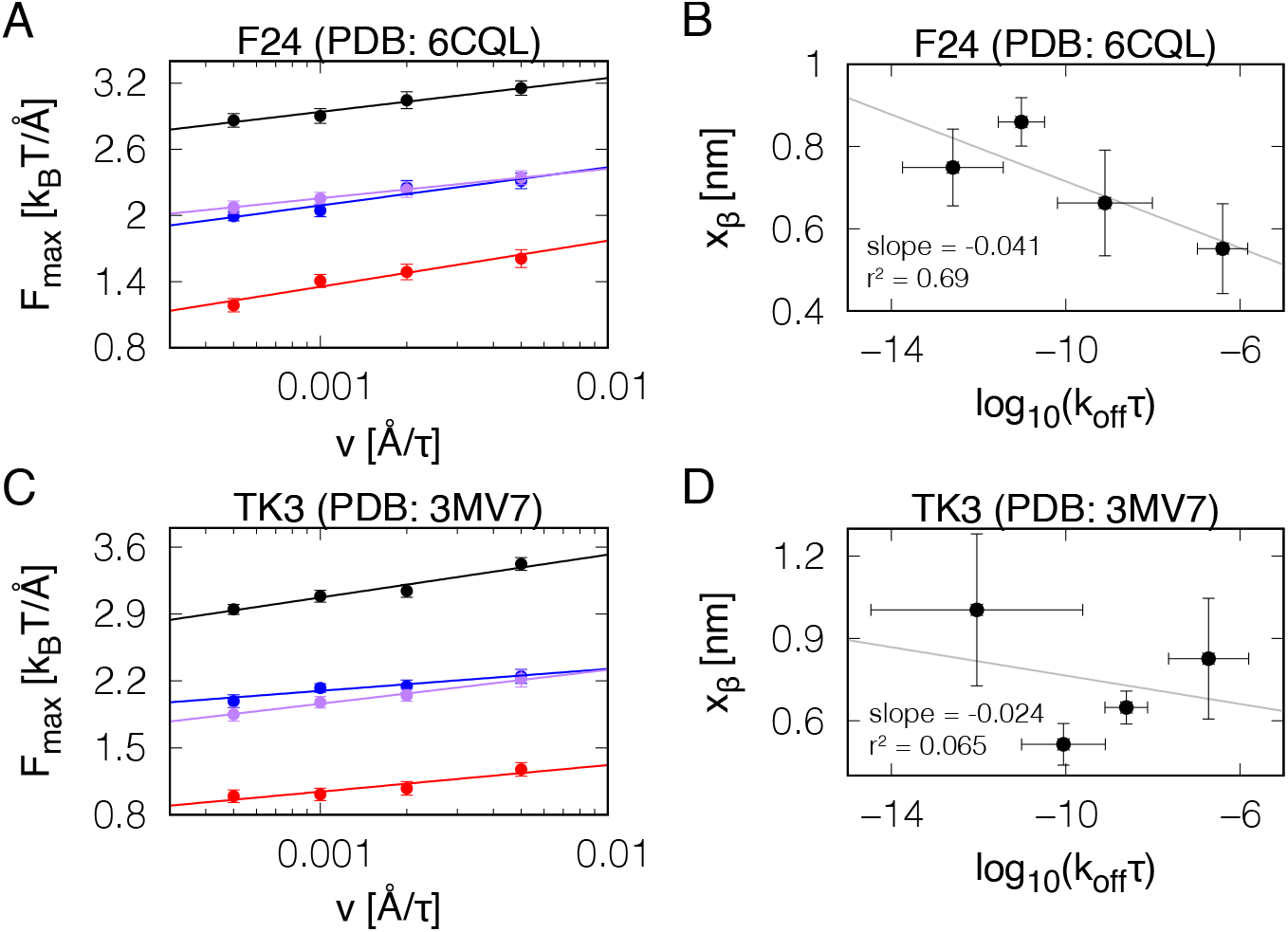
Negative x_β_ k_off_ correlation is reproduced in two other TCRs. Analogous to Figure 3D–G but for two other pMHCs in complexes with the F24 TCR **(A–B)** and the TK3 TCR **(C–D)**. The negative correlation between x_β_ and log(*k*_off_) is observed also for these TCR-pMHC complexes (B and D).

**Figure S6.**
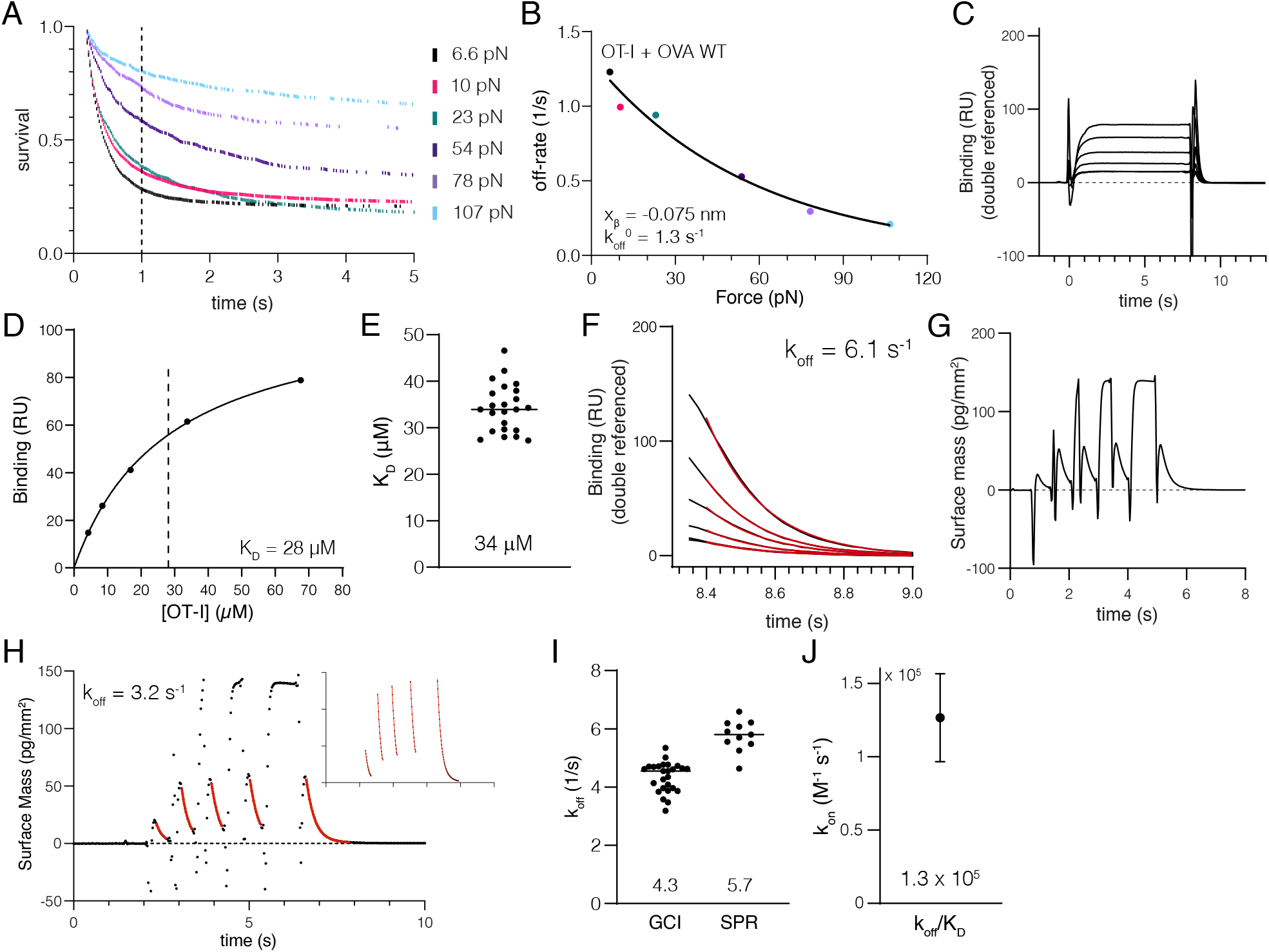
The OT-I TCR binds OVA with low affinity and a catch bond. (A) Example of bead survival over time at different velocities/forces. The time to measure survival (1 s) is shown as dotted line. **(B)** Fitting the off-rates at different forces to Bell’s model. Free parameters are the zero-force unbinding rate and the force sensitivity (*x*_*β*_). Colors correspond to forces in (B). The number of independent experiments that were combined to produce the estimated off-rates are: 11 (OT-I/OVA) **(C)** Representative surface plasmon resonance (SPR) sensogram of OT-I TCR binding immobilized OVA pMHC at five different concentrations of TCR. **(D)** Steady-state RU from (A) plotted over the TCR concentration fitted by a one-site specific binding model to determine K_D_ (solid line). **(E)** Fitted K_D_ values from multiple experiments (N=23). **(F)** Dissociation phase from (A) fitted with a one phase exponential decay model to determine *k*_off_ (solid red lines). **(G)** Representative grating-coupled interferometry (GCI) sensogram of OT-I TCR binding immobilized OVA pMHC at one concentration of TCR injected for increasing durations. **(H)** Fit of the entire sensogram data from (G) determined *k*_off_ (red line). **(I)** Fitted *k*_off_ values from four and two independent experiments with different TCR and/or pMHC concentrations using GCI (N=25) or SPR (N=11), respectively. **(J)** Calculated on-rate (*k*_on_) using the fitted values of *k*_off_ (4.3 s^*−*1^) and K_D_ (34 *µ*M).

